# Hunting strategies and the lateral line of *Astyanax mexicanus* larva

**DOI:** 10.1101/791418

**Authors:** Marina Yoffe, Kush Patel, Eric Palia, Samuel Kolawole, Amy Streets, Gal Haspel, Daphne Soares

**Affiliations:** Biological Sciences, New Jersey Institute of Technology, Newark NJ 07102; The University of North Carolina at Chapel Hill, NC 27514; Westfield High School, Westfield, NJ 07090; Queensland Brain Institute University of Queensland St. Lucia, QLD 4072

**Keywords:** Lateral line, adaptation, fish, caves, preadaptation, evolution

## Abstract

The lateral line is the primary modality fish use to create a hydrodynamic image of their environment. These images contribute to a variety of behaviors, from rheotaxis to escape responses. Here we discern the contributions of visual and lateral line modalities in hunting behavior of larvae that have developed under different photic conditions. In particular, cave animals have a hypertrophied sense of mechanosensation, and we studied the common animal model cavefish *Astyanax mexicanus* and its closest related surface relative. We raised larvae in a diurnal light-dark regimen and in complete darkness. We then examined the distribution of neuromasts in their lateral lines, and their hunting performance in light and dark conditions, with and without the contribution of the lateral line. We report that all larva depend on the lateral line for success in hunting and that surface fish raised in the dark have a greater dependency on the lateral line,

## Introduction

It is well known that troglobitic animals have enhanced mechanosensation as a compensatory mechanism for the loss of vision (Niemiller and Soares, 2013; Soares and Niemiller, 2018). In fish mechanosensation is mostly sensed through a series of receptors on the skin, collectively called the lateral line. Fishes’ lateral lines are composed of individual end organs called neuromasts. In turn, neuromasts are composed of hair cells. These cells have stereocilia which extrude into the surrounding water from the skin. Neuromasts are present in two different forms, as single organs on the surface of the skin and in canals. Both function by encoding the deflection of a gelatinous cupula that surrounds the stereocilia of the hair cells. Each type is sensitive to a particular bandwidth of frequencies, creating a complex hydrodynamic image of the surrounding aquatic environment in the brain of the fish (Coombs et al., 2000; Montgomery et al. 2001). When the lateral line has been ablated, fish show impaired detection of prey (Liang et al., 1998; Coombs 1999; Pohlmann et al., 2004), evasion of predators (Blaxter and Fuiman, 1989), avoidance of obstacles (Abdel-Latif et al., 1990; Hassan et al., 1992), mating (Satou et al., 1994), schooling (Partridge and Pitcher, 1980; Partridge 1981), and rheotaxis (Montgomery et al., 1997; Baker and Montgomery, 1999, Suli et al. 2012). Blind adult cavefish of the genus *Astyanax* show a hypertrophy of the lateral line (Teyke, 1990; Montgomery et al. 2001) and a distinct lateral line centric behavior called the vibration attraction behavior (Yoshizawa et al. 2014).

Hunting in larval fish has been extensively studied (Hunter, 1980; Hubbs and Blaxter, 1986). Zebrafish for example are highly visual hunters (Bianco et al. 2011), while other fish larvae rely on other sensory modalities for survival (Gerlach et al., 2007; Wright et al., 2005). It is clear that prey movement is an important cue for fish larva to hunt. The brine shrimp hatchling (*Artemia salina*) create disturbanes in the water by using a rowing propulsion to move via a pair of trunk limbs. Fish larvae in the laboratory are often fed *Artemia* that are approximately 0.5 mm in size. Such prey swim and 1.8 mm/s (Williams, 1994) and live in a low Reynolds number environment. Fish larvae therefore, have to detect microscale turbulence with their lateral lines in order to use mechanosensation to capture prey.

*Astyanax mexicanus* has various cavefish populations that have independently evolved in a world of darkness. It is difficult to imagine how they survive and maneuver as larvae in the dark waters of the caves; especially because there are seasonal floods and food is scarce. In contrast, the closest living relative population of *Astyanax*, surface fish that lives in local rivers, have visual-centric behaviors, having large eyes even as larva. Cavefish larva on the other hand have small eyes that degenerate during the early stages of life (Jeffery and Martasian, 1998; Yamamoto and Jeffery, 2000). These two forms of *Astyanax* are closely related, having diverged as early as 20K years ago (Funey et al., 2018), and can still interbreed in laboratory setting, being considered one species. Here we set out to examine the lateral line and hunting behavior of the larva of *Astyanax* because it might give us an insight into why the genus is so successful at colonizing the underground environment. In this study, we examined emergence of neuromasts and hunting during the first two weeks of development of the lateral line system in a representative *Astyanax* cave lineage (Pachón) and compared with the surface fish.

## Methods

### Larval care

Larvae from Pachón cave lineage as well as surface fish larvae were raised at 23 °C under two light conditions. One group was raised in the dark incubator with minimal exposure to light, while the other group was held in an incubator with 12:12h light /dark conditions. Larvae were fed live *Artemia* daily starting 6 days post-fertilization. Animal husbandry and experimentation were covered under the Rutgers University IACUC protocol #201702685.

### Hunting

Experiments were conducted on 7, 11, and 14 days old larvae under light or dark conditions. A day prior to the experiment, larvae were fed live *Artemia* to satiation and then were transferred into a treatment (fish water containing 0.002% gentamicin) or control (fish water) petri dishes. Preceding the experiment, 30 minutes adaptation time in hunting illumination conditions was given to the fish after their transfer into 6-well experimental plates. Hunting illumination conditions were either dark, in which plates were kept in a dark cabinet, or lit, in which they were kept on top of the same cabinet with ambient room light. Each well received ten live *Artemia* (prey) for the duration of one hour, after which larvae were removed from the wells and the remaining *Artemia* were counted.

### Imaging of the Lateral Line

Fish larvae at 7, 11, and 14 days post fertilization were treated for 3 min with 0.05% DASPEI (2-4- dimethylaminostyryl-N- ethylpyridinium iodide) solution (Sigma Aldrich). DASPEI stain highlights specific mitochondria in the intact neuromasts of the lateral line (Van Trump et al., 2010). Following staining, an epifluorescence stereozoom microscope with a phototube was used to collect images of larvae under a band-pass GFP filter set (excitation 450-490 nm, emission 500-560 nm). Using the microscope, images were processed of both the lateral and dorsal views, so that the number of cranial, trunk, and total neuromasts could be observed. Neuromasts were then quantified using the free FIJI (ImageJ) software. Data visualization for figures was conducted on R platform (R Core Team).

### Statistics

Comparison of data from different groups was done by 3-way ANOVA on Matlab (using the functions anovan and multcompare, our code is available upon request). This allowed evaluation of the effect of each factor and their interactions and the testing for significant difference between each pair of groups with multiple comparisons post hoc (Tukey’s honest significant difference HDC test).

## Results

We compared the number of *Artemia* hunted in 60 min by eight groups of surface fish larvae that were raised in dark or 12:12 light conditions, were treated with gentamicin or control fish water, and hunt in lit or dark conditions (Figure 1). We used a three-way ANOVA to test the null hypotheses that there is no difference in the population means for different treatments and their interaction, and found that each of the treatments caused a significant difference: larvae raised in the dark caught less *Artemia* than those raised in 12:12 light conditions (F=6.4, p=0.0116), those treated with gentamicin caught less *Artemia* (F=588, p=10^-77) and those hunting in the dark caught less *Artemia* (F=123, p=10^-24). Interactions between factors were also significant, except from the interaction between raising and hunting conditions. In general, larvae with intact lateral lines hunted successfully under all conditions (multiple comparisons post-hoc, Tukey’s honest significant difference HSD; 9.3±0.3 9.7±0.3 8.9±0.3 8.6±0.3, raised in dark or 12:12 conditions, and hunting in light or dark respectively), while all larvae with ablated lateral line hunted poorly in the dark (0.9±0.3 1.7±0.3, raised in dark or 12:12 conditions respectively). The exceptions are larvae with ablated lateral lines that hunted in the light. Under those conditions, larvae that were raised in the dark (4.9±0.3) caught less artemia (p=0.007) than those raised in 12:12 light conditions (6.4±0.3).

**Figure 1.**
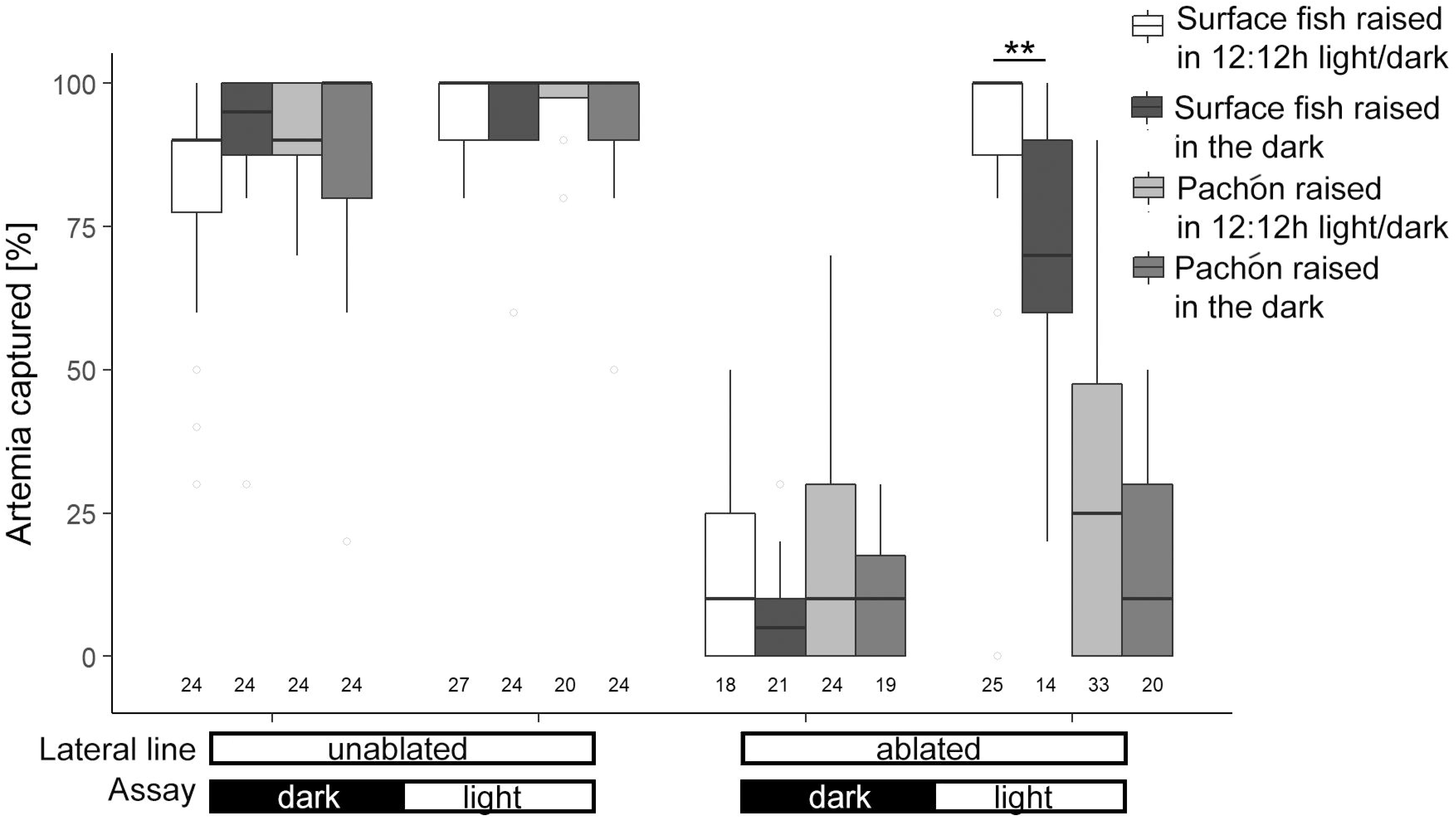
Dark raised surface fish larvae rely on their lateral line for prey capture more than those raised under 12:12h conditions. Prey capture assay of Pachón and surface fish larvae raised in the dark or under light /dark conditions in four different scenarios: gentamicin treated in larval performance in the dark (lateral line ablated raised in the dark) or in the light (lateral line ablated raised in the 12:12h), and controls in the dark (raised in the dark) and in the light (raised in the light). Numbers below each column represent the sample size of the column.

With similar analysis of Pachón cavefish larvae we found that each of the treatments caused a significant difference: larvae raised in the dark caught less *Artemia* than those raised in 12:12 light conditions (F=6.13, p=0.0143), those treated with gentamicin caught less *Artemia* (F=743, p=10^-63) and those hunting in the dark caught less *Artemia* (F=8.61, p=0.004). Interactions between factors were not significant. In general, larvae with intact lateral lines hunted successfully under all light conditions (multiple comparisons post-hoc, Tukey’s honest significant difference HSD; 9.2±0.3 9.7±0.3 8.7±0.3 8.9±0.3, raised in dark or 12:12 conditions, and hunting in light or dark respectively), while all larvae with ablated lateral line hunted poorly in dark or lit conditions (1.7±0.3 2.9±0.4 1.1±0.4 1.9±0.4, raised in dark or 12:12 conditions, and hunting in light or dark respectively). In contrast to surface fish, raising conditions did not affect hunting success of Pachón larvae.

We compared the number of neuromasts on the head and trunk of larvae among twelve groups: 7, 11, or 14 days old, surface fish or Pachón cavefish larvae, and raised in dark or 12:12 light conditions (Figure 2). We used three-way ANOVA to test the null hypotheses that there is no difference in the population means for different ages, fish type, raising conditions, and their interaction. For cranial neuromasts there were significant differences in the number of neuromasts between Pachón cavefish and surface fish (F=31.7, p=5×10^-8) and among 7, 11, and 14 day old larvae (F=66.4, p=10^-23), but not between larvae raised in different light conditions (F=0.02, p=0.88); also significant were the differences along interactions between types and age or raising conditions (F=16.3, p=2×10^-7; F=3.8, p=0.05), but not the interaction between age and raising conditions (F=0.17, p=0.84). In general, 14 days old Pachón cavefish larvae had more cranial neuromasts than all other groups (multiple comparisons post-hoc, Tukey’s honest significant difference HSD; 42.0±0.42 42.4±0.6, raised in dark or 12:12 conditions, respectively), while all other larvae were not different from each other (35.0±0.5 37.3±0.5 37.8±0.6 38.2±0.6, in 7 and 11 days old Pachón, raised in dark and 12:12 light conditions, and 35.6±0.5 36.7±0.5 38.2±0.4 35.1±0.5 36.2±0.6 37.3±0.6, in 7, 11, and 14 days old surface fish, raised in dark and 12:12 light conditions).

**Figure 2.**
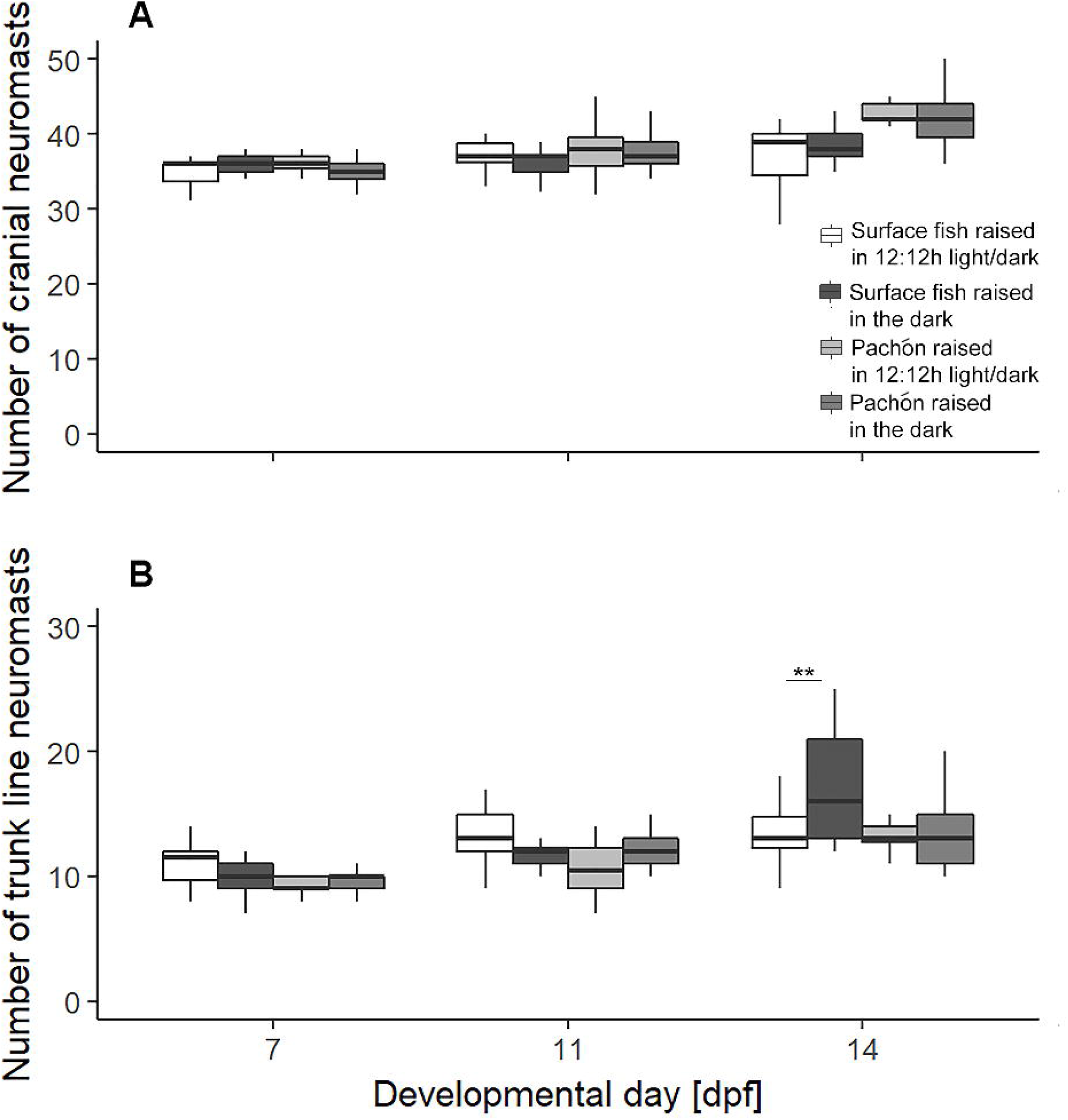
Surface fish raised in the dark have more neuromasts in the body than 12:12h light/dark raised surface fish. A) Neuromast count on one side of the cranium. B) Neumast count of one side of the trunk lateral line.

For trunk lateral line there were significant differences in the number of neuromasts between Pachón cavefish and surface fish (F=18.3, p=3×10^-5) and among 7, 11, and 14 day old larvae (F=59.4, p=9^-22), but not between larvae raised in different light conditions (F=2.33, p=0.13); also significant were the differences along interactions between age and fish type or raising conditions (F=3.1, p=0.05, F=7.8 p=0.0005), but not the interaction between fish type and raising conditions (F=0.09, p=0.76). In general, older larvae had more trunk neuromasts. Comparing larvae of different groups that were raised in dark or 12:12 light conditions, only 14 day old surface fish that were raised in the dark (16.6±0.4) had more neuromasts than those raised in 12:12 light conditions (14.3±0.5; p=0.007; multiple comparisons post-hoc, Tukey’s honest significant difference HSD), while all other pairs of larval groups were not different from each other (9.2±0.5 11.3±0.5 14.1±0.4 10.0±0.6 11.5±0.6 11.9±0.6, in 7, 11 and 14 days old Pachón, raised in dark and 12:12 light conditions, and 10.1±0.5 12.4±0.5 10.6±0.5 12.4±0.6, in 7 and 11 days old surface fish, raised in dark and 12:12 light conditions).

## Discussion

### Main findings

Our experimental design was formulated to discern the contributions of visual and mechanosensory aspects of hunting in *Astyanax* larvae that have developed in different photic environments. Results show that all larvae are successful in hunting prey using the lateral line. In the dark, surface fish perform as well as cavefish, suggesting that they are using sensory modalities other than vision. Only fish in which the neuromasts were ablated showed a decrease in hunting performance. Surface fish raised in the 12:12 light regimen showed an almost perfect performance when tested in the light with mechanosensory lateral line ablated. Suggesting that this behavior is mediated by vision. The same is not true for dark-raised surface fish with ablated lateral line. These fish performed worse, suggesting that during their raising period in the dark they have adjusted their behavior to hunt relying on mechanosensation. Furthermore surface fishes that were raised in the dark had more trunk lateral line neuromasts than when raised in the 12:12 regimen. An alternative explanation to our results is that the larvae that hunt more simply move more and have an increased chance of randomly encountering prey. However, in larval fish, ingesting prey is an outcome of not just encountering but also attacking prey (Dower et al. 1997). The prey capture that occurs in ablated fish in the dark could also be partially mediated by chemosensory detection, which is consistent across all testing conditions.

We found that presence and location of neuromasts were most consistent in the head, with very little variation. These neuromasts are under evolutionary pressure because they are responsible for the vibration attraction behavior (Yoshizawa et al. 2014) in cavefish and likely prey capture in surface and cave fish. We saw variation in the body neuromasts that have not been previously reported to be used in prey capture.

### Surface fish preadaptation?

A long standing question has been why is *Astyanax* the only group that has been successful in invading caves in the Sierra del Abra, which rivers host many other species? With our results we can propose that surface *Astyanax* have an advantage when born to live in dark caves because the development of neuromasts seems to be plastic and influenced by the visual environment. We would like to argue that their plastic dependency in mechanosensation has allowed the first generation of surface fish larvae to survive a novel dark environment.

## Supporting information

hunting

## Grant support

NIH R15EY027112

The data that support the findings of this study are available from the corresponding author upon reasonable request.

## Supplementary Material

Table 1. The total number of neuromasts on one side of the fish at different developmental points.

Figure S1. The hunting performance of larvae at A) 7 and B) 14 days post fertilization. Prey capture assay of Pachón and surface fish larvae raised in the dark or under light /dark conditions in four different scenarios: gentamicin treated in larval performance in the dark (lateral line ablated raised in the dark) or in the light (lateral line ablated raised in the 12:12h), and controls in the dark (raised in the dark) and in the light (raised in the light). Numbers below each column represent the sample size of the column.

