## Supplementary figures and images for "Hunting strategies and the lateral line of *Astyanax mexicanus* larva"

### hunting

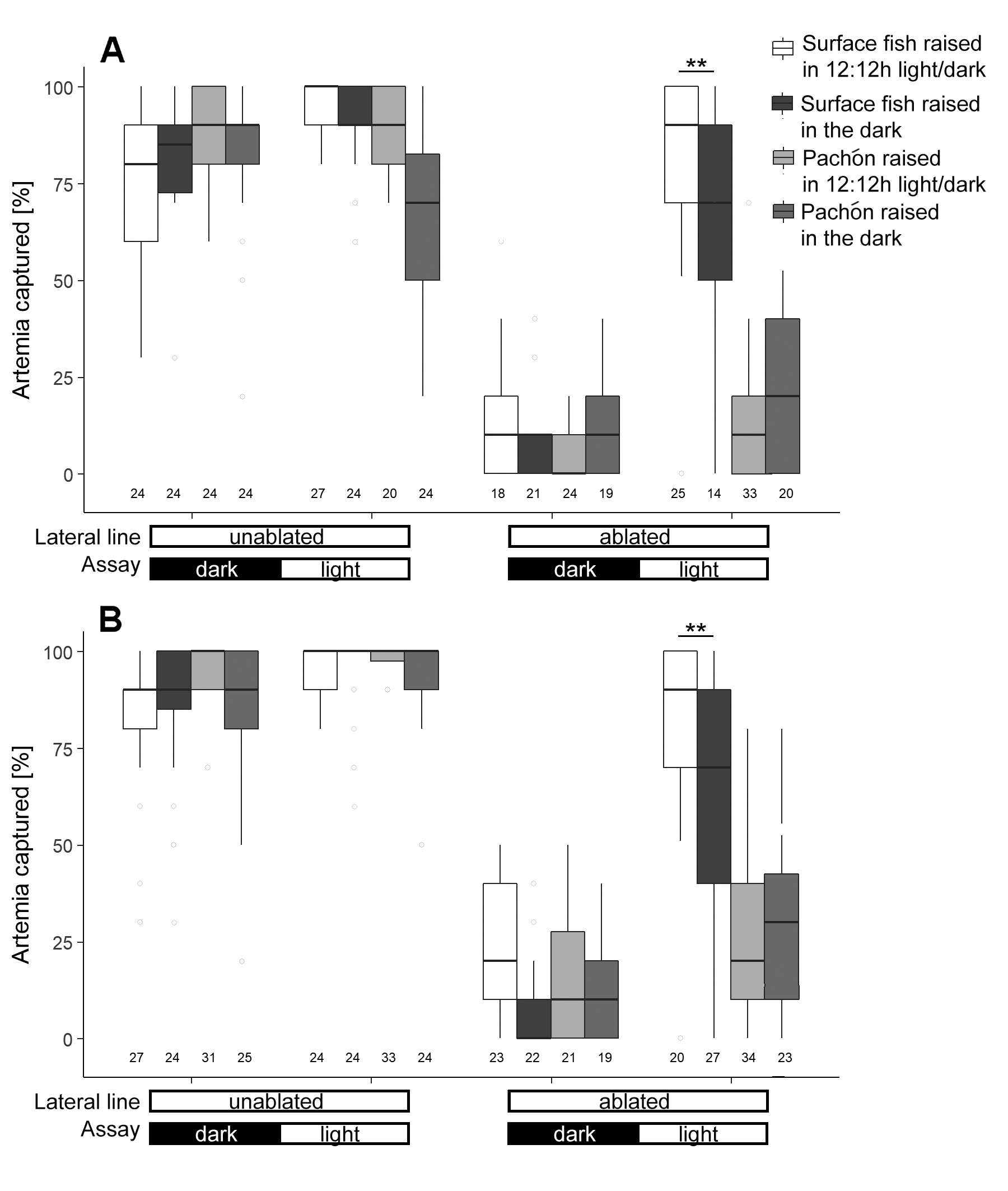

### table 1

**Table 1.** The total number of neuromasts on one side of the fish at different developmental points.

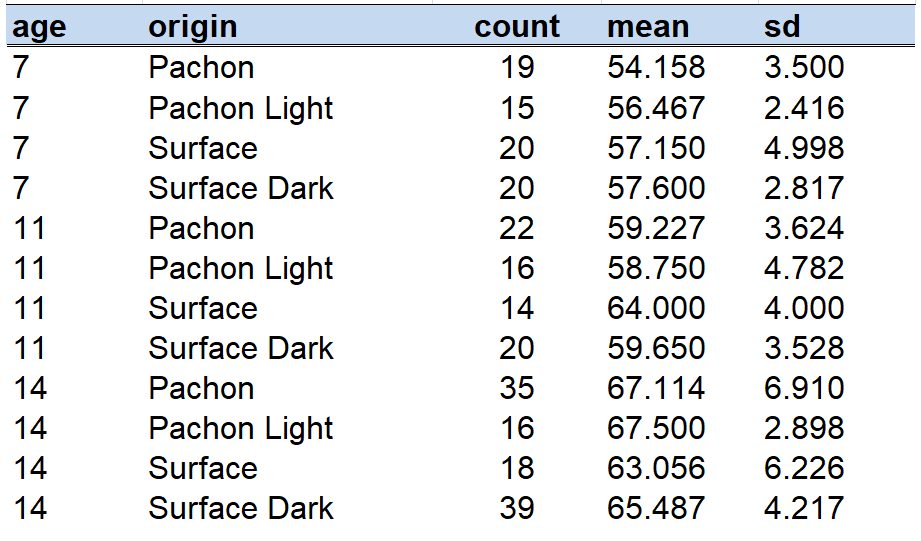
